# Allometry, sexual selection, and evolutionary lines of least resistance shaped the evolution of exaggerated sexual traits within the genus *Tyrannus*

**DOI:** 10.1101/2021.07.09.451836

**Authors:** MN Fasanelli, P Milla Carmona, IM Soto, DT Tuero

**Affiliations:** Instituto de Ecología, Genética y Evolución de Buenos Aires - IEGEBA (CONICET-UBA), Departamento de Ecología, Genética y Evolución -DEGE, Facultad de Ciencias Exactas y Naturales, Universidad de Buenos Aires, Int. Guiraldes 2160, Buenos Aires, Argentina; Laboratorio de Biología Integral de Sistemas Evolutivos. DEGE, Facultad de Ciencias Exactas y Naturales, Universidad de Buenos Aires, Int. Guiraldes 2160, Buenos Aires, Argentina; Laboratorio de Ecosistemas Marinos Fósiles, Instituto de Estudios Andinos Don Pablo Groeber (CONICET-UBA), Intendente Güiraldes 2160, Ciudad Universitaria, C1428EHA Buenos Aires, Argentina

**Keywords:** constraints, morphology, birds, tails, diversification

## Abstract

Variational properties hold a fundamental role in shaping biological evolution, exerting control over the magnitude and direction of evolutionary change elicited by microevolutionary processes that sort variation, such as selection or drift. We studied the Tyrannus genus, as a model for examining the conditions and drivers that facilitate the repeated evolution of exaggerated, secondary sexual traits in the face of significant functional limitations. We study the role of allometry, sexual selection, and their interaction on the diversification of tail morphology in the genus, assessing whether and how they promoted or constrained phenotypic evolution. The exaggerated and functionally-constrained long feathers of deep-forked species, *T. savana* and *T. forficatus*, independently diverged from the rest of the genus following the same direction of main interspecific variation common to the entire cluster of species. However, at a macroevolutionary scale those axes summarising both sexual dimorphism and allometric variation of the deep-forked species were aligned with the between-species maximum variation axis of non deep-forked species. Thus, we are presenting evidence of amplified divergence via the co-option and reorientation of allometric shape variation involved in a sexual selection process that repeatedly drove morphology along a historically favoured direction of cladogenetic evolution.

## Introduction

The direction and magnitude of evolutionary change are largely controlled by evolutionary processes that sort variation such as natural selection and drift [1,2], but also by the sources of that variation, such as the genetic architecture underlying the affected phenotypes [3]. Intrapopulational variation *per se* is a major evolutionary factor that can enhance or impede response to selection by modulating the amount of raw material available for phenotypic changes elicited by other processes [4,5]. Over short evolutionary timescales, selective responses may be biased by strong genetic correlations or simply by the absence of genetic variation thus precluding or deviating evolutionary change towards certain phenotypes [6–8]. This includes the early stages of between-species divergence, where evolution is predicted to be biased along “genetic lines of least resistance” defined by the additive genetic variance-covariance matrix “*G”* [4,9].

However, natural selection has its agency at the organismal level, acting upon available phenotypic variation [1]. As the organismal phenotype is the result of developmental processes unfolding during ontogeny, the latter can favour the emergence of particular phenotypes (e.g. [10]), and thus has the potential to establish preferred directions for long-term evolution [11]. In this sense, they constitute “phenotypic lines of least resistance” defined by P, the phenotypic variance-covariance matrix (see [12]). The influence of these constraints is well documented for relatively short evolutionary timescales (i.e., less than 2 million years; [4,13–18];see [19] for a discussion).

Nonetheless, the influence of genetic and developmental constraints over greater evolutionary timescales is yet to be determined [1,20] as empirical studies are scarce (but see [19,21]). Genetic constraints are frequently disregarded when studying species diversification or other long-term evolutionary phenomena under the argument that the genetic architecture of trait covariation may evolve as well [22–24] consequently altering potential genetic lines of least resistance [25]. Selection and drift can alter the G matrix [23,26–28] but the signature of genetic constraint may persist over large timescales even in the presence of an evolving genetic architecture [19].The study of phenotypic lines of least resistance can greatly improve the study of developmental constraints, as it is both easier to estimate than and highly correlated to their genetic counterpart, as well as highly informative in regard to evolution (e.g., [12,29]).

Allometry is a classic example of trait covariation and a central issue in the literature addressing developmental integration, as changes in many morphological, physiological and life history traits are highly correlated with changes in organ or body size [30–32]. Because ontogenetic and evolutionary allometric relationships are often evident over large size ranges, and since allometric slopes (both static and ontogenetic) are usually rather conserved among closely related species, allometry has been considered as a constraint for morphological evolution [10,33,34]. In this view, the existence of allometric patterns is often interpreted as evidence of strong ontogenetic, physiological or other biological mechanisms that restrain or limit the rate and direction of evolution and diversification [35,36]. An alternative view is to consider the processes underlying the observed allometric patterns as facilitators of evolution or an adaptation on itself [30,37–40]. Under this perspective, which is often discussed in the context of classic studies ([30]; see a discussion in [41]), the effect of selective processes that shape trait relationships or phenotypic responses are amplified by the existent covariance structure.

This latter view is widespread in ornithological studies, as allometric patterns are considered a major factor contributing to phenotypic variation across a range of avian clades [42,43]. Previous studies focused on avian traits (beaks, tails and wings) showed how ecological factors drive the evolution of those structures with the aid of allometry [44–46]. For example, Bright et al. [47] found a strong covariation between skull shape and size, showing that exploitation of allometric relationships is an effective mechanism by which raptor birds may modify their feeding ecology.

*Tyrannus* (Lacépède 1799) is a New World genus of passerines comprising thirteen species with a common ancestor about 5 MY before present and with a widespread American distribution, from Canada to Argentina [44,48,49]. Those species inhabit a wide range of environments, such as savannahs, woods and even urban areas [50,51] Some species are seasonal migrants, although migratory behaviours within the genus are variable (e.g., partial, short-distance and long distance migration; [49,52,53]). They prey primarily on flying insects and forage almost exclusively during flight, although they also feed from high perches or on the ground [54–56]. The group includes two deepfork-tailed (hereafter, DF) species, the Scissor-tailed (*T. forficatus*) and the Fork-tailed (*T. savana*) Flycatchers, from North America and South America, respectively (Figure 1, [51,57,58]).

**Figure 1.**
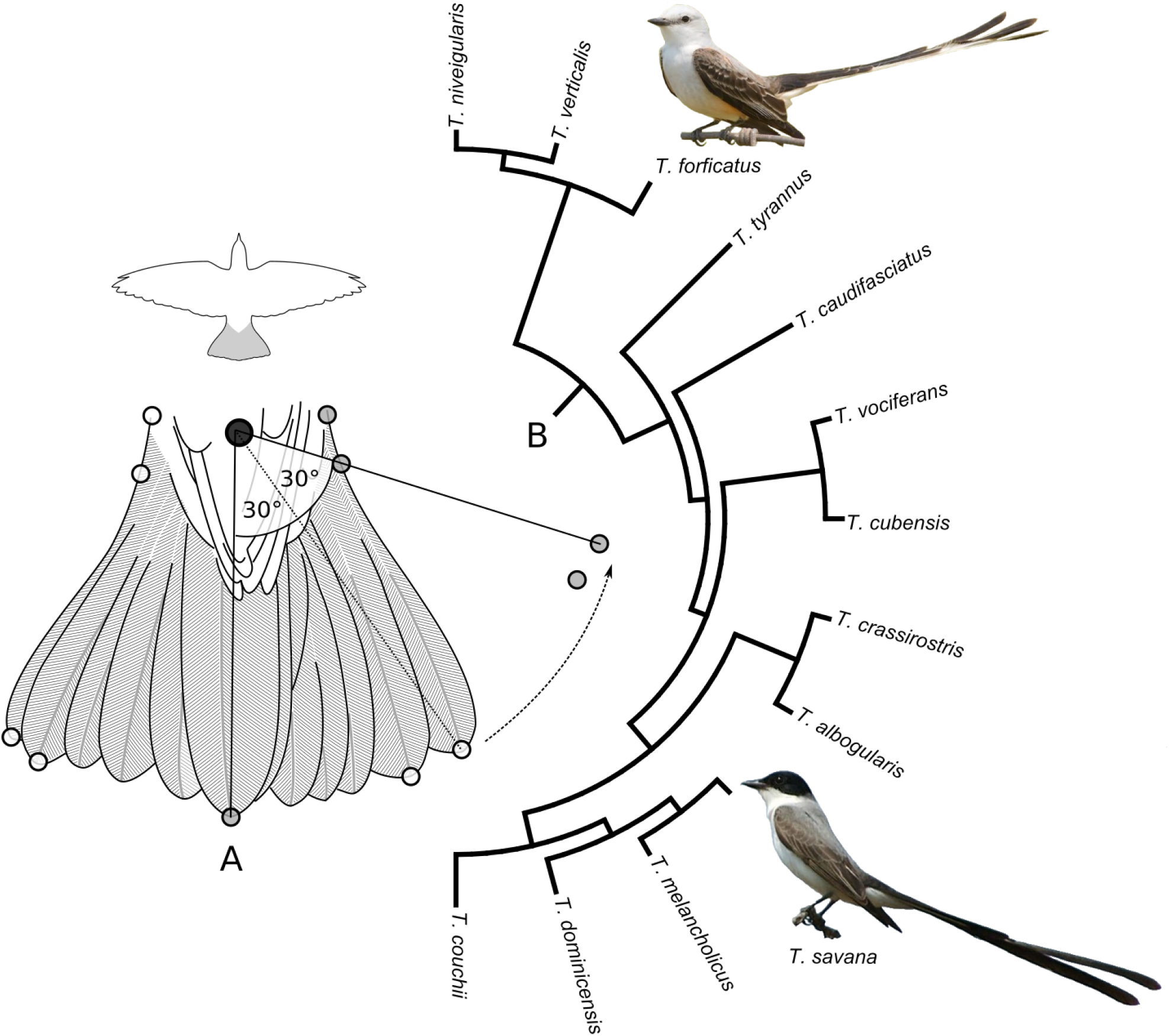
A) Schematic representation of a bird tail, showing the original and post-rotated position of the nine landmarks (semi-open circles), as well as the centroid of the tail base (larger closed circle) used as pivot; only right-sided landmarks (semi-open gray circles) were retained for further analysis. B) Cladogram of the *Tyrannus* species [48], with the two deep-fork tailed species depicted.

Derived from independent monophyletic lineages [59], these DF species present disjunct geographic distributions [60] and are characterised as socially monogamous, with both male and female involved in parental care [61]. Both species show extremely elongated tails in males and females, as well as a significant sexual dimorphism in tail length attributed to the action of sexual selection [58,62,63].

Here, we address the genus *Tyrannus* as a model for examining the conditions and drivers that facilitate the repeated evolution of exaggerated, secondary sexual traits in the face of significant functional limitations, with particular focus on the role of allometry, sexual selection, and their covariation, on the diversification of tail morphology in the genus. In particular, we estimate and explore the relationship between two major sources of intraspecific variation (sexual dimorphism and allometric variation) and the main directions of macroevolutionary change that accrued during the evolution of the clade, assessing whether they promoted or otherwise constrained the evolution of tail morphology by establishing phenotypic lines of least resistance.

## Materials and methods

### Data collection

The thirteen species of *Tyrannus* were represented in a sample of 281 museum specimens (range: 12-39 specimens per species, average: 21.6 specimens per species; American Museum of Natural History, AMNH, USA and Museo Argentino de Ciencias Naturales, MACN, Argentina). Only adult skins of both sexes with rectrices in good condition were included.

Tails were photographed in dorsal view from a standardised vertical plane. Tail morphology was captured by placing nine landmarks on the image of the open tail of each specimen: 1) the tips of the central rectrix, as well as #2–#5) the two outermost rectrices, #6–#7) the pygostyle insertion point, and #8–#9) the inflection point formed by the opened outer rectrix at the base of the tail of each side (Figure 1).

Due to the fragile feathers of museum material, tails were only opened at an angle of 60° (Figure 1A). Since an opening of 120° is the standard in studies of bird tail morphology [66], the two landmarks marking the tip of the outermost rectrices of each side were further rotated 30°, using the centroid of landmarks #6–#9 (defining the ‘base’ of the tail) as pivot. In order to avoid problems reported elsewhere for supervised versions of Principal Component Analysis (PCA) and its algebraic equivalents (i.e., between-group PCA and Partial Least Squares; e.g.,[64,65]), the four landmarks corresponding to the left side of the tail (having served to determine the central axis of symmetry) were discarded and all subsequent analyses were carried out using the five remaining landmarks (Figure 1A; note that these landmarks were mirrored to represent tails in all figures).

Centroid size (CS) of each landmark configuration was calculated to be used as a proxy for tail size, after which the sample of landmark configurations was subjected to Generalized Procrustes Analysis and projected into tangent space [66], thus separating size from shape variation to be analysed independently.

Body size was computed for a subset of 260 specimens for which measures of wing, bill and tarsus lengths, as well as body weight, were available. A PCA of these variables was performed and the scores of specimens along the first axis were used as a body size proxy. The relationship between tail and body size was assessed by performing Linear Regressions, using raw and sexually-corrected size estimates. The latter ‘correction’ was performed within each species by computing theresiduals from a linear model of each size estimate on sex. Tail and body sizes were z-standardised in all cases to make values comparable.

### Interspecific shape variation

Raw shape variation within each species was processed to remove intraspecific variation by computing the shape residuals resulting from a Linear Model of tail shape on tail size and sex. This ‘corrected’ data set was further refined by means of between-group PCA (bgPCA; see [67] and references therein), performed using all *Tyrannus* species but *T. savana*and *T. forficatus* as *a priori* groups. These procedures were applied to avoid the heavy influence that deep-forked shapes (and to a lesser extent, the intraspecific shape variation) exert over the orientation of the axes resulting from a regular PCA. Instead, a series of synthetic shape variables (bgPCs), rotated to maximize interspecific variation among those species possessing nondeep-forked (hereafter, NDF) tail morphologies, were constructed (bgPC1 being renamed from here onward as the between-species maximum variation axis, or BMV axis). Only then, samples and mean shapes belonging to *T. savana* and *T. forficatus* were projected into the resulting bgPCs. By using the interspecific variation displayed by NDF species as the basis for constructing our shape variables, we sought to enhance interpretability of our results by establishing the main directions of evolutionary variation resulting from ‘normal’ within-clade dynamics as reference.

### Intraspecific shape variation

Starting again from the raw shape variables, four types of intraspecific shape variation were isolated within each species. First, the axis of maximum variation (from now on, within-species maximum variation axis, or WMV axis) was estimated through the first PC resulting from a PCA of raw shape variation. Second, an axis representing allometric shape variation was constructed, using Partial Least Squares (PLS) of shape on log-transformed centroid size (in this context, this procedure finds a linear combination of shape variables maximising the correlation with changes in size; see [68]) and computing the shapes expected under the estimated relationship by sampling the range of realised PLS scores at regular intervals. Third, an axis representing differences between sexes was generated by computing male and female mean shapes along the axis resulting from a new bgPCA using sex as grouping factor. Fourth, an axis representing residual variation (i.e., not explained by changes in either size, sex or their interaction) was obtained by performing a PCA over the corresponding shape residuals. These axes were characterised using their eigenvector’s coefficients and depicted by projecting their expected shapes into the morphospace.

Statistical significance of shape differences attributable to intersexual and allometric variation was assessed for each species separately through Procrustes Distances-based Linear Models (using sex and log-transformed centroid size as independent variables, respectively). The magnitude of this difference was measured using the partial Procrustes distance implied in each shape transformation. Whether the BMV and the allometric and intersexes axes are more aligned than expected by chance was tested separately for each species by comparing the Pearson correlation between the corresponding eigenvector coefficients to the distribution of the correlation between randomly permuted eigenvector coefficients, generated through 999 iterations [32].

### Macroevolutionary pattern of divergence

A molecular phylogeny was obtained from *birdtree.org* and *onezoom.org* from Jetz et al. [48,59] and Hackett et al. [69] in order to provide phylogenetic structure for the analyses (Figure 1B). Both tail shapes and patterns of intraspecific variation were reconstructed for hypothetical ancestors by optimising the species’ mean shape configurations and sets of corresponding eigenvectors within a maximum parsimony framework [70]. This methodology reconstructs ancestral traits maximising similitude accounted for common ancestry. Phylogenetic structure was included in morphospaces by projection of shape configurations corresponding to the tips and nodes of the phylogeny, along with their genealogical relationships, to create a phylomorphospace [71].

Finally, in order to explore whether between-species divergence took place along lines of least resistance, as well as the nature of those lines, the alignment between axes was tested for every possible triad of an ancestor and its two descendants, using the same permutation test approach described above. These comparisons correspond to a) the alignment between the inter-descendants axis (i.e., the bgPC separating the pair of descendants’ shapes) and the ancestral WMV, and b) the alignment between the ancestral WMV and ancestral allometric, intersexes, and residual axes of variation.

All analyses were conducted in the R environment [72] using the packages Morpho [73], geomorph [74], shapes [75], abind [76], retistruct [77], spdep [78], ape [79] and phytools [80]. The only exceptions were the digitisation of tail landmarks, carried out using tpsDig [81] and the ancestral node reconstruction, implemented in Tree Analysis using New Technology (TNT; [82]).

## Results

### Size and shape

Raw body and tail sizes displayed a significant positive correlation in four out of the 13 species of *Tyrannus: T. forficatus*, *T. niveigularis*, *T. savana* and *T. tyrannus*. However, this correlation was lost in the two deep-fork-tailed species when the sexually-corrected sizes were used instead (Figure 2, Supplementary Table 1).

**Figure 2.**
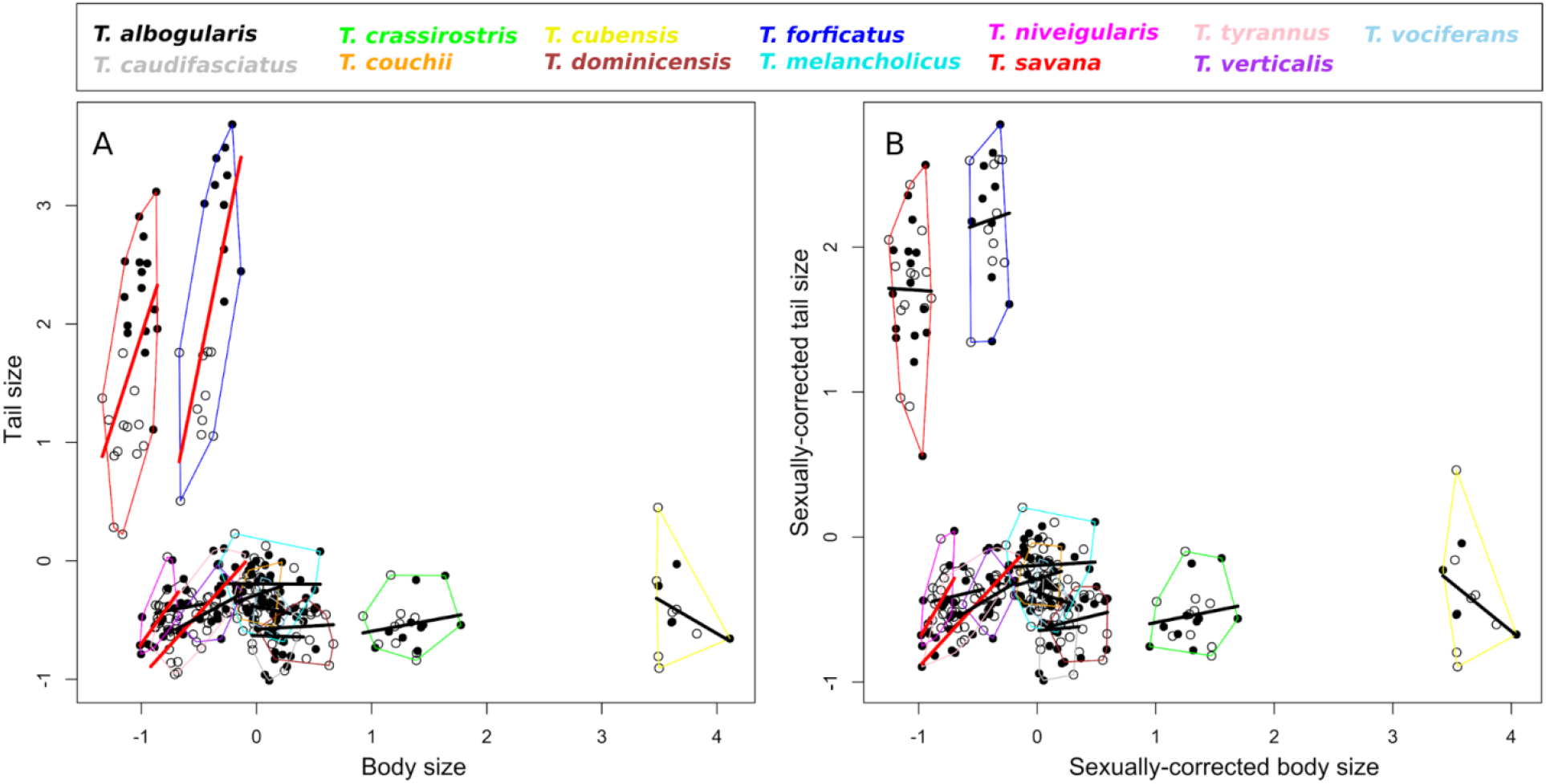
Z-standardized raw (A) and sexually-corrected (B) tail size versus body size plots. Open circles represent females, whereas closed circles represent males. Solid thick lines are linear regression trend lines; Red lines indicate slopes significantly different from 0.

The first three Principal Components resulting from bgPCA accounted for 98.91% of the corresponding variance (Figure 3A–D, Supplementary Figure 1). Variation along the first bgPC(BMV, capturing 72.86% of the original shape variance) describes differences in the degree of forking of the tail and relative base width, with negative values representing forked tails with outermost rectrices slightly more elongated that inner outer rectrices, and positive values representing rounded tail fans. The second bgPC (20.95% of total variance explained) on the other hand captures differences in tail elongation, tail base width and relative positions of the tips of outermost rectrices relative to the sagittal plane. Finally, bgPC3 (5.11% of the original shape variance) represents subtle differences in separation of outermost rectrices’ tips. Most *Tyrannus* species display an overlapped distribution in the resulting phylomorphospace. Unsurprisingly, *T. savana* and *T. forficatus* are the exception, diverging markedly from the rest of the genus along bgPC1, and from each other along bgPC3 (Figure 3A–B).

**Figure 3.**
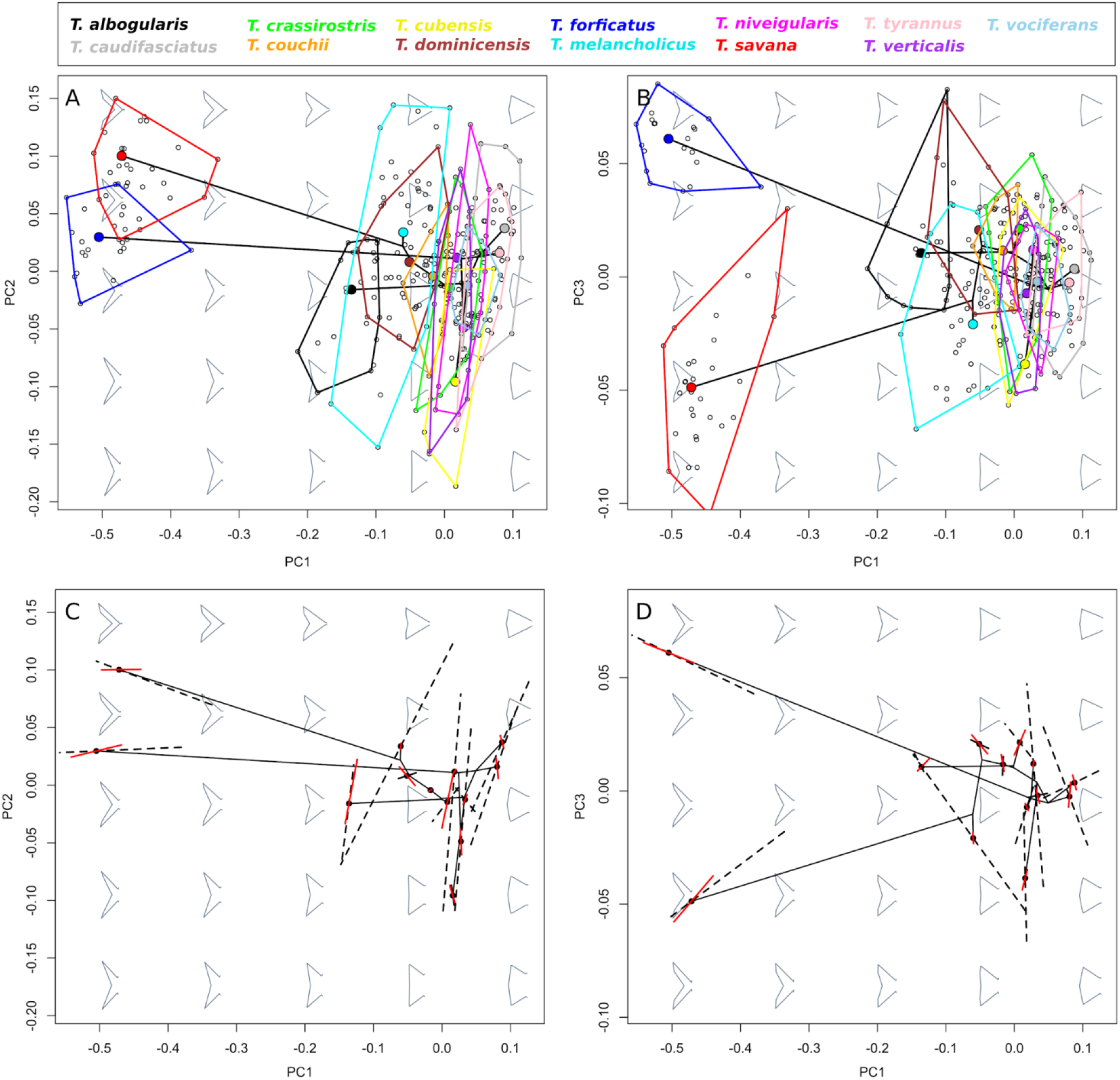
A–B) Phylomorphospaces formed by the first three bgPCs displaying the distribution of the full sample of allometrically- and sexually-corrected specimens (open circles) and species’ mean shapes (coloured closed circles). C–D) Intraspecific axes of morphological variation (allometric variation: black dashed lines; intersexes variation: solid red lines) projected into phylomorphospace and centered around each species’ mean shape.

### Inter- and intraspecific shape variation

In general, *Tyrannus* species with NDF tails displayed a reduced allometric and intersexes variation (with the exception of sexual dimorphism in *T. albogularis*), as evidenced by both the magnitude and statistical significance of the implied shape transformations (Table 1). These axes are significantly aligned with each other in four species (Table 2). However, although their alignment with the BMV axis differ (Figure 3C–D), none of these species display a significant correlation between the latter and allometric or intersexes axes (the sole exception being the intersexes axis of *T. dominicensis*; Table 2, Figure 3C–D).

**Table 1.** Procrustes distances implied in shape transformations for intraspecific allometric and intersexes variation, along with the residual degrees of freedom, F values and p-values from Procrustes Linear Models. Significant allometric patterns or sexual dimorphism are highlighted in bold.

|  | Allometric variation | Sexual dimorphism |
| --- | --- | --- |
| *T. albogularis* | 0.044 (df = 14; F = 0.48; p = 0.679) | **0.059(df = 14; F = 3.14; p = 0.042)** |
| *T. caudifasciatus* | 0.065 (df = 19; F = 1.24; p = 0.242) | 0.015 (df = 19; F = 0.21; p = 0.890) |
| *T. couchii* | 0.042 (df = 15; F = 0.77; p = 0.470) | 0.017 (df = 15; F = 0.43; p = 0.739) |
| *T. crassirostris* | 0.037 (df = 18; F = 0.41; p = 0.682) | 0.052 (df = 18; F = 3.05; p = 0.076) |
| *T. cubensis* | 0.089 (df = 10; F = 1.21; p = 0.290) | 0.021 (df = 10; F = 0.22; p = 0.873) |
| *T. dominicensis* | 0.035 (df = 19; F = 0.55; p = 0.645) | 0.031 (df = 19; F = 1.15; p = 0.319) |
| *T. forficatus* | **0.116 (df = 18; F = 15.26; p > 0.001)** | **0.064 (df = 18; F = 13.44; p > 0.001)** |
| *T. melancholicus* | 0.039 (df = 37; F = 0.31; p = 0.759) | 0.010 (df = 37; F = 0.13; p = 0.928) |
| *T. niveigularis* | 0.069 (df = 17; F = 1.12; p = 0.275) | 0.023 (df = 17; F = 0.45; p = 0.608) |
| *T. savana* | **0.165 (df = 27; F = 20.28; p > 0.001)** | **0.070 (df = 27; F = 10.95; p > 0.001)** |
| *T. tyrannus* | 0.075 (df = 24; F = 3.02; p = 0.06) | 0.023 (df = 24; F = 0.78; p = 0.427) |
| *T. verticalis* | 0.038 (df = 19; F = 0.36; p = 0.718) | 0.011 (df = 19; F = 0.14; p = 0.943) |
| *T. vociferans* | 0.028 (df = 18; F = 0.71; p = 0.519) | 0.013 (df = 18; F = 0.52; p = 0.729) |

**Table 2.** Pearson correlations between the allometric and intersexes axes of each species and the BMV, along with the p-value from permutation tests. Statistically significant results are highlighted in bold.

|  | Allometric – Intersexes | Allometric – Interspecific | Intersexes – Interspecific |
| --- | --- | --- | --- |
| *T. albogularis* | **0.677 (p = 0.028)** | 0.081(p = 0.654) | 0.232(p = 0.407) |
| *T. caudifasciatus* | 0.517(p = 0.123) | 0.377(p = 0.181) | -0.391(p = 0.371) |
| *T. couchii* | **0.761(p = 0.008)** | -0.032(p = 0.971) | 0.199(p = 0.495) |
| *T. crassirostris* | 0.190(p = 0.603) | -0.492(p = 0.171) | -0.226(p = 0.348) |
| *T. cubensis* | -0.334(p = 0.344) | 0.026(p = 0.123) | 0.421(p = 0.322) |
| *T. dominicensis* | 0.508(p = 0.153) | -0.382(p = 0.337) | **-0.793(p = 0.017)** |
| *T. forficatus* | **0.987(p > 0.001)** | **-0.98(p > 0.001)** | **-0.97(p > 0.001)** |
| *T. melancholicus* | 0.451(p = 0.191) | -0.500(p = 0.084) | -0.047(p = 0.804) |
| *T. niveigularis* | **0.897(p = 0.002)** | -0.087(p = 0.712) | 0.117(p = 0.860) |
| *T. savana* | **0.974(p > 0.001)** | **-0.666(p = 0.038)** | **-0.656(p = 0.034)** |
| *T. tyrannus* | **0.810(p = 0.004)** | 0.282(p = 0.291) | -0.208(p = 0.690) |
| *T. verticalis* | 0.154(p = 0.672) | 0.048(p = 0.700) | 0.419(p = 0.244) |
| *T. vociferans* | 0.339(p = 0.339) | -0.497(p = 0.210) | 0.368(p = 0.211) |

The situation changes dramatically for the DF species, as both *T. savana* and *T. forficatus* exhibit significant allometric and intersexes variation (Table 1). However, the most remarkable aspect of these species is the alignment of both allometric and intersexes axes of variation not only with each other, but with the main BMV axis of the NDF *Tyrannus* species (i.e., they are approximately parallel; Table 2, Figure 3C–D).

As for the ancestor-descendants comparisons, most of the correlation between the different reconstructed types of intraspecific variation and the reconstructed ancestral WMVaxis can be attributed to residual variation (Table 3). In this regard, the main difference between comparisons excluding and focused on DF species is an increased contribution (roughly 50%) of allometric variation for the latter in relation to the former. The pattern of contribution of the different reconstructed types of intraspecific variation to the correlation with the reconstructed inter-descendants axis was very similar for comparisons involving only NDF tailed species. In contrast, intersexes variation accounted for most of the correlation between the reconstructed ancestral WMV and the reconstructed inter-descendants axis of variation in comparisons involving only DF species (Table 3).

**Table 3.** Accumulated correlations between ancestral axes of intraspecific variation and the ancestral WMV (left) or the interdescendants axis (right), associated with each ancestor-descendants comparison. Values are relativised to the highest value of each column.

|  | vs. ancestral axis<br>of major<br>intraspecific<br>variation |  |  | vs. major axis of<br>inter-descendant<br>variation |  |
| --- | --- | --- | --- | --- | --- |
|  | Regular clade<br>contrasts | Deep-fork species<br>related contrasts |  | Regular clade<br>contrasts | Deep-fork species<br>related contrasts |
| Residual | 1 | 1 |  | 1 | 0.199 |
| Allometric | 0.548 | 0.810 |  | 0.623 | 0.484 |
| Intersexual | 0.611 | 0.588 |  | 0.540 | 1 |

## Discussion

Allometric patterns are important aspects contributing to diversification of avian morphology [42,43], in concert with ecological pressures on phenotypes [44–47]. Sexual selection is also an important evolutionary force in the context of avian evolution, affecting behaviour, colouration and morphology of birds [83,84].

In the context of bird tail morphology and sexual selection, this topic has received considerable attention [38,40,42] being generally addressed from the ‘classic’ Huxley-Jolicoeur school of allometry, i.e., by studying the relation between tail and body size [85]. The general consensus emerging from these studies is that secondary sexual traits with exaggerated morphologies will tend to show positive allometry (i.e., larger tails will be associated with larger bodies) as the result of the balance between metabolic costs and reproductive advantage. Our findings support this prediction, as both *T. savana* and *T. forficatus* show positive tail-to-body size allometry (Figure 2, Suplementary Table 1).

However, these results represent only the first of two very different allometric phenomena this study is concerned with. The second, and the one that is the main focus of this work, is the relation between trait size and shape (framed within the ‘modern’ Mosimann-Gould approach to allometry; [85]). A very straightforward takeaway from our results is that this kind of allometric variation and sexual dimorphism are both amplified and evolutionary coupled in the two independently evolved *Tyrannus* species with exaggerated tail morphologies (Figures 2–3). Ongoing sexual selection has been reported in both *T. savana* and *T. forficatus* [58,62]. Still, the coupling of sexual dimorphism (a natural by-product of sexual selection [86,87] with allometric shape variation suggests that this powerful agent of evolutionary change is being fed with a pervasive and ecologically significant source of morphological variation. This is in line with the notion that sexual selection has played a central role in the evolution of exaggerated tail morphologies in birds [88–91], but also shines light on why such disparate shapes have been able to be realised in the first place.

Nonetheless, there are several other *Tyrannus* species showing both coupled patterns of intersexual and allometric shape variation and NDF tails (Table 2). The missing piece lies in another result: the marked morphological differences between *T. savana* and *T. forficatus* and the rest of the genus can be summarised as a monotonic divergence along the same direction of main interspecific variation extracted from the cluster of species with NDF tails (the BMV). But perhaps the most striking finding of this study is that, at a macroevolutionary scale, axes summarising both sexual dimorphism and allometric variation of *T. savana* and *T. forficatus* are aligned with the BMV of NDF species.

This latter axis can be considered as resulting from the accumulation of cladogenetic change (under the reasonable assumption of the prevalence of non-directional anagenetic patterns of change; [92]), and will therefore reflect the directions of phenotypic evolution favoured during between-species divergence associated to regular within-clade dynamics in *Tyrannus*. This divergence follows the direction of maximum intraspecific variation present in the ancestor –mainly corresponding to residual intraspecific variation, with modest contributions of sexual dimorphism and allometry– for the majority of the cladogenetic events involving the evolution of NDF species (Table 3). Thus, regular within-clade phenotypic evolution can be regarded as following a prevailing phenotypic line of least resistance [4]. In speciation events leading to a DF species, however, the overall contribution of allometry to ancestral intraspecific variation is substantially higher. Moreover, interdescendants divergence correlates strongly with ancestral sexual dimorphism, hinting the action of sexual selection (Table 3). Putting all this information together, we suggest that two conditions are necessary for the evolution of exaggerated tails in *Tyrannus*: 1) the co-option of allometric shape variation as the primary source of raw material for sexual selection, and 2) the alignment of the latter process with a macroevolutionary line of least resistance, resulting in fast divergence through an open lane of macroevolutionary change.

If these lessons can be generalised, we should expect other cases of sexually-related exaggerated or highly elaborated traits to represent an amplification of ‘regular’ within-clade dynamics, catalysed by a process of sexual selection and fuelled by allometric –or perhaps other substantial intraspecific source of– morphological variation. In particular, we predict that 1) species with exaggerated or elaborated morphologies will diverge along the main axis of interspecific variation already present in the cluster of related species with regular morphology, and 2) the latter axis will be aligned with the axes describing intraspecific differences between sexes and allometric (or other kinds of) morphological variation.

Ornithological literature has openly understood the design of bird tail as a flight device, not only from the historically-vast observational registers, but most probably through the widely used theoretical models of bird flight. A ‘functionalist’ approach has been heavily supported when addressing the evolution of the many bird species showing marked sexual dimorphism and elaborated tail morphologies (e.g., [86,87,91]), with most of the literature revolving around the relative contribution and putative antagonism of natural and sexual selection. In this context, net selection on tail’s morphology is conceived as the compromise between its role as secondary sexual trait –i.e., its effects on mating success– and as a structure involved in flight –i.e., its effects on aerodynamic performance– [89,93], invoking both biomechanical conflicts [94](Evans 2004) and physiological trade-offs [95,96].

In contrast, the other side of the functionalist coin, namely the ‘structuralist’ aspects (i.e., general biases in the production of variation that introduce limits, but also channels, for evolution; see [1]) of this study case, have received much less attention. The existence of variation, long recognised as the water to selection’s mill, has been generally treated as something that is given (or not) for tail form without further consideration. Even allometry, a topic subject to recent debate, is treated as an expression of the tension between natural and sexual selection (see above). Yet, the present study hints that the existence of some highly elaborated morphologies (a feature strongly associated with selection) crucially depends on the interaction of different types, levels and specific properties of variation. Our intention is not to argue against the importance of selection (a process our results are consistent with), but to vindicate the creative role that evolutionary changes in direction, magnitude and coupling of different sources of variation (whatever their particular mechanistic base might be) plays into reaching a satisfactory explanation. In this particular case, the co-option (and potential reorientation) of allometric shape variation as a source of raw material for processes of sexual selection driving mean morphology along a historically favoured direction of cladogenetic evolution resulted in amplified divergence.Evolutionary changes in the properties of variation, as well as their feedback with selection and other processes, need not to follow the same evolutionary dynamics than the current selective regimes identified and discussed so far in the literature, and represent an open avenue for future research.

## Acknowledgments

We are grateful to the curators, technicians and collection managers of the bird divisions of The American Museum of Natural History (George F. Barrowclough, Joel L. Cracraft and Paul Sweet) and the Museo Argentino de Ciencias Naturales “Bernardino Rivadavia” (Darío Lijtmaer and Yolanda Davies) for access to materials under their care and for their kind assistance. MNF was a recipient of a Collection Study Grant from The Frank M. Chapman Memorial Fund of the AMNH Museum.

## Authors contributions

**MNF** collected the primary data, performed the morphometric quantification, participated in data analysis and in the design of the study and critically revised the original draft.

**PMC** conceived and participated in the conceptualization and design of the study, carried out the data analysis and visualisation and drafted the manuscript along with IMS.

**IMS** conceived the study and contributed to the study conceptualization and design, coordinated the study, provided funding, contributed to the data analysis and statistics and drafted the manuscript along with PMC.

**DTT** conceived and coordinated the study, participated in the design of the study, provided funding and critically revised the original draft.

All authors performed revisions and editing of the initial draft and gave final approval for publication and agreed to be held accountable for the work performed therein.

---

**Supplementary Table 1.** Results from regressions of tail size on body size for both raw and sexually corrected estimates of size. Statistically significant results are highlighted in bold.

|  | Raw |  |  |  |  | Sexually-corrected |  |  |  |
| --- | --- | --- | --- | --- | --- | --- | --- | --- | --- |
|  | Residual<br>df | F<br>values | p-values | Slopes |  | Residual<br>df | F<br>values | p-values | Slopes |
| *T. albogularis* | 14 | 0,437 | 0,52 | 0,181 |  | 14 | 0,557 | 0,468 | 0,196 |
| *T. caudifasciatus* | 19 | 0,006 | 0,939 | -0,031 |  | 19 | 0,03 | 0,865 | 0,079 |
| *T. couchii* | 12 | 0,842 | 0,377 | 0,337 |  | 12 | 0,544 | 0,475 | 0,252 |
| *T. crassirostris* | 18 | 0,553 | 0,467 | 0,179 |  | 18 | 0,373 | 0,549 | 0,162 |
| *T. cubensis* | 10 | 0,86 | 0,375 | -0,567 |  | 10 | 1,074 | 0,325 | -0,663 |
| *T. dominicensis* | 19 | 0,047 | 0,831 | 0,057 |  | 19 | 0,486 | 0,494 | 0,201 |
| *T. forficatus* | **18** | **16,004** | **0,001** | 4,793 |  | 18 | 0,067 | 0,799 | 0,29 |
| *T. melancholicus* | 36 | 0,001 | 0,973 | -0,006 |  | 36 | 0,064 | 0,801 | 0,047 |
| *T. niveigularis* | **16** | **8,399** | **0,01** | 1,368 |  | **16** | **7,613** | **0,014** | 1,403 |
| *T. savana* | **27** | **9,112** | **0,005** | 3,025 |  | 27 | 0,004 | 0,951 | -0,055 |
| *T. tyrannus* | **21** | **28,512** | **>0,001** | 1,073 |  | **21** | **34,599** | **>0,001** | 0,892 |
| *T. verticalis* | 18 | 3,598 | 0,074 | 0,525 |  | 18 | 2,067 | 0,168 | 0,486 |
| *T. vociferans* | 15 | 0,208 | 0,655 | -0,191 |  | 15 | 0,411 | 0,531 | -0,285 |

**Supplementary Figure S1.**
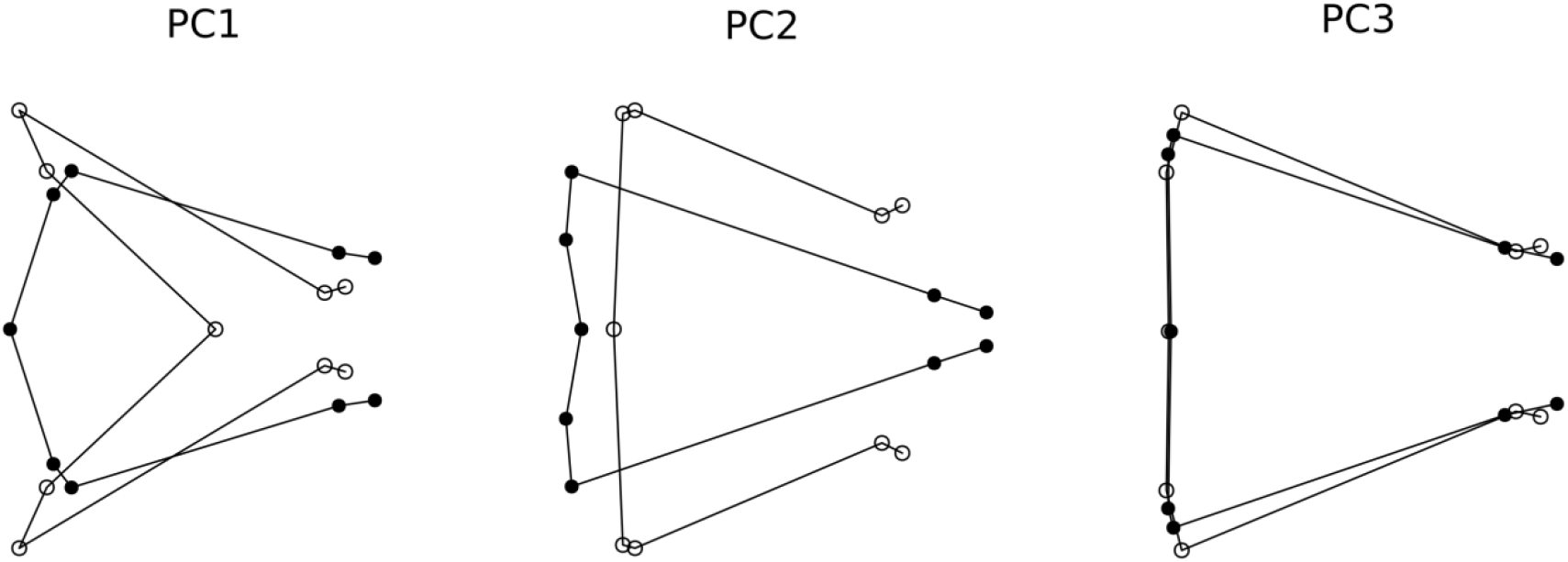
Shape transformations represented by the first three Principal Components resulting from PCA of Procrustes shape coordinates of NDF *Tyrannus* species. Open circles are used for landmarks of the configuration representing negative extremes, whereas closed circles are used for positive extremes.

